# Human melanoma cell lines that possess wild-type *BRAF* alleles but are dependent on *ERBB4* and *ERBB2*

**DOI:** 10.1101/2024.08.22.609260

**Authors:** Vipasha Dwivedi, Lauren M. Lucas, Rees Cooke, Jeniffer Davis, Ella Wilson, Markelle Scott, Kaitlyn O’Daniel, Chloe Dion, Jaxon Kerley, Madison Zelan, Nicholas DeFeo, Victoria Huffman, Madison N. Ingrao, David J. Riese

## Abstract

Metastatic skin cutaneous melanomas that contain wild-type *BRAF* alleles typically possess an activating mutation in a *RAS* allele or a loss-of-function mutation in an *NF1* allele (“*BRAF*-WT&*RAS*/*NF1-*mutant melanomas”). Nonetheless, these tumors remain a significant clinical challenge; they are resistant to MEK and BRAF inhibitors, their response to immune checkpoint inhibitors is less robust than the response of *BRAF* mutant melanomas to these agents, and additional validated targets for therapeutic intervention have yet to be identified.

Previous work from our laboratory has demonstrated that *ERBB4* is required for the proliferation of the IPC-298, MEL-JUSO, MeWo, and SK-MEL-2 *BRAF*-WT&*RAS*/*NF1-*mutant melanoma cell lines. Surprisingly, the synthetic constitutively dimerized and active Q646C *ERBB4* mutant allele appears to strongly inhibits the proliferation of *BRAF*-WT&*RAS*/*NF1-* mutant melanoma cell lines. Given that we have also previously demonstrated that ERBB4-ERBB2 and ERBB4-EGFR heterodimers are more potent drivers of proliferation than are ERBB4 homodimers, here we begin to test the hypothesis that ERBB4 heterodimers drive the proliferation of *BRAF*-WT&*RAS*/*NF1-*mutant melanoma cell lines.

Here we demonstrate that the kinase-deficient (dominant-negative) *ERBB2* K753A mutant allele inhibits the clonogenic proliferation of the IPC-298, MEL-JUSO, and MeWo *ERBB4*-dependent, *BRAF*-WT&*RAS*/*NF1-*mutant melanoma cell lines. Moreover, the kinase-deficient (dominant-negative) *EGFR* K721A mutant allele inhibits the clonogenic proliferation of the MeWo cell line, but not the IPC-298 or MEL-JUSO cell lines. Finally, the clonogenic proliferation of the SK-MEL-2 *ERBB4*-dependent, *BRAF*-WT&*RAS*/*NF1-*mutant melanoma cell line is unaffected by the *ERBB2* K753A or *EGFR* K721A dominant-negative mutant alleles. We discuss these findings in the context of our hypothesis that ERBB4 heterodimers drive the proliferation of *BRAF*-WT&*RAS*/*NF1-*mutant melanoma cell lines.

## INTRODUCTION

Melanoma is a form of skin cancer that is particularly aggressive and deadly. Fortunately, 50% of melanomas harbor a gain-of-function mutation in a *BRAF* allele and these tumors are particularly responsive to immune checkpoint inhibitors (ICIs) or a combination of BRAF and MEK inhibitors [1, 2]. Most melanomas that possess wild-type *BRAF* alleles harbor a gain-of-function mutation in a *RAS* allele or a loss-of-function mutation in an *NF1* allele. Nonetheless, no targeted therapeutics (including MEK inhibitors) are effective against these *BRAF* WT, *RAS/NF1* mutant (“*BRAF*-WT&*RAS/NF1*-Mut”) melanomas, and ICIs are less effective against these tumors than against *BRAF* mutant melanomas [1-3]. Thus, we have sought targets in *BRAF*-WT&*RAS/NF1*-Mut melanomas that cooperate with the *RAS/NF1* mutant alleles to drive these tumors.

A combination of *in silico* and *in vitro* analyses from my laboratory suggest that elevated signaling of wild-type *ERBB4* [1] or mutational activation of *ERBB4* [4] appears to drive the proliferation of the IPC-298, MEL-JUSO, MeWo, and SK-MEL-2 *BRAF*-WT&*RAS*/*NF1-*mutant melanoma cell lines.

ERBB4 (HER4) is a member of the ERBB family of receptor tyrosine kinases (RTKs), which includes the epidermal growth factor receptor (EGFR), ERBB2 (HER2/Neu), and ERBB3 (HER3). ERBB4 possesses extracellular ligand-binding domains, a single-pass hydrophobic transmembrane domain, an intracellular tyrosine kinase domain, and intracellular tyrosine residues that function as phosphorylation sites **(Figure 1)**. Ligand binding to EGFR, ERBB3, or ERBB4 stabilizes the receptor extracellular domains in an open conformation competent for symmetrical homodimerization and heterodimerization of the receptor extracellular domains. The dimerization of the extracellular domains enables asymmetrical dimerization of the receptor cytoplasmic domains. Phosphorylation of one receptor monomer on tyrosine residues by the tyrosine kinase domain of the other receptor monomer (“cross-phosphorylation”) ensues **(Figure 2)**. This tyrosine phosphorylation creates binding sites for effector proteins and activation of downstream signaling pathways [5].

**Figure 1.**
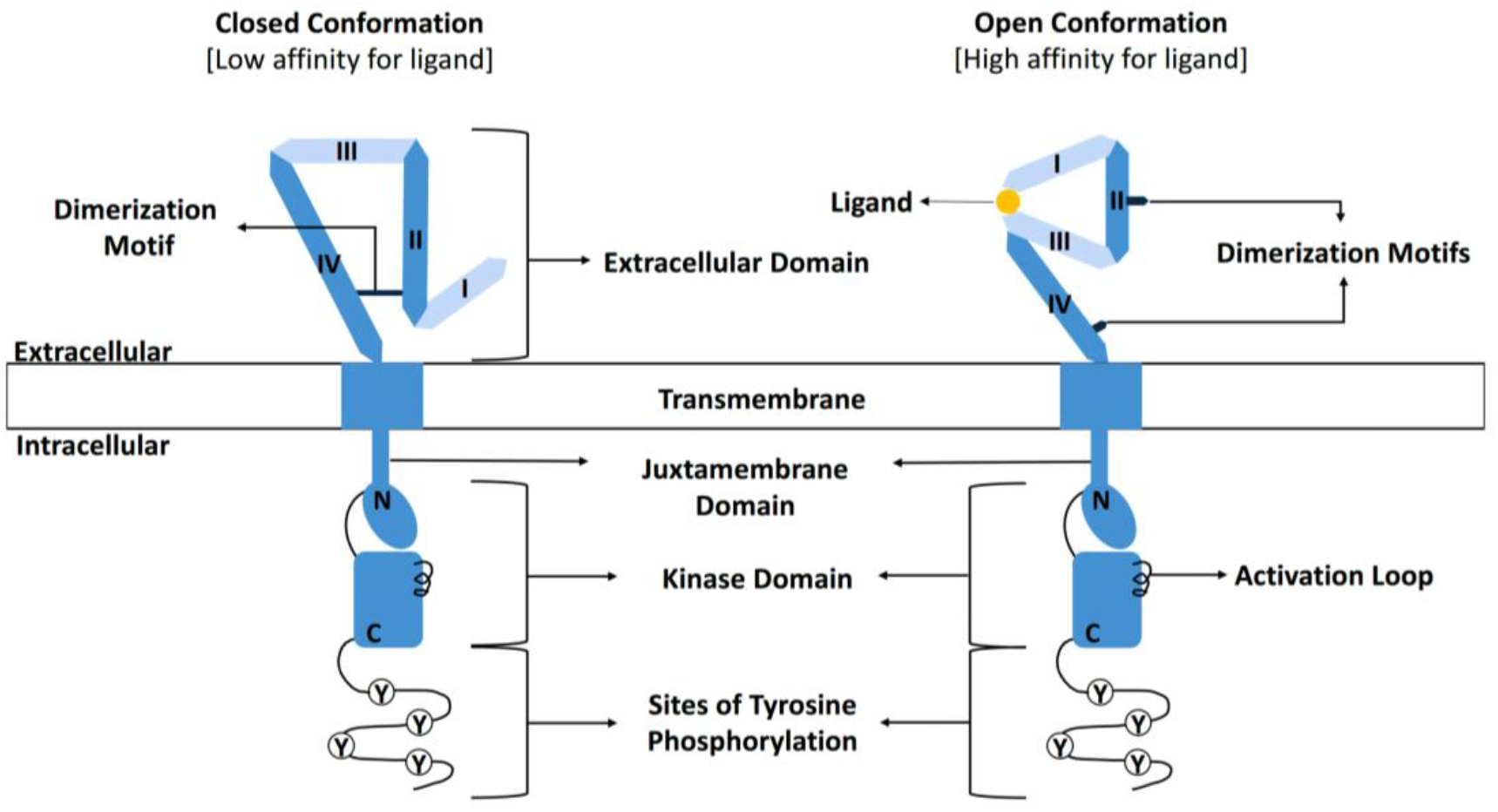
The extracellular domains of ERBB4 exist in an equilibrium between the closed conformation that has low affinity for ligand and buried dimerization motifs and the open conformation that has high affinity for ligand and has exposed dimerization motifs. Adapted from [4, 5].

**Figure 2.**
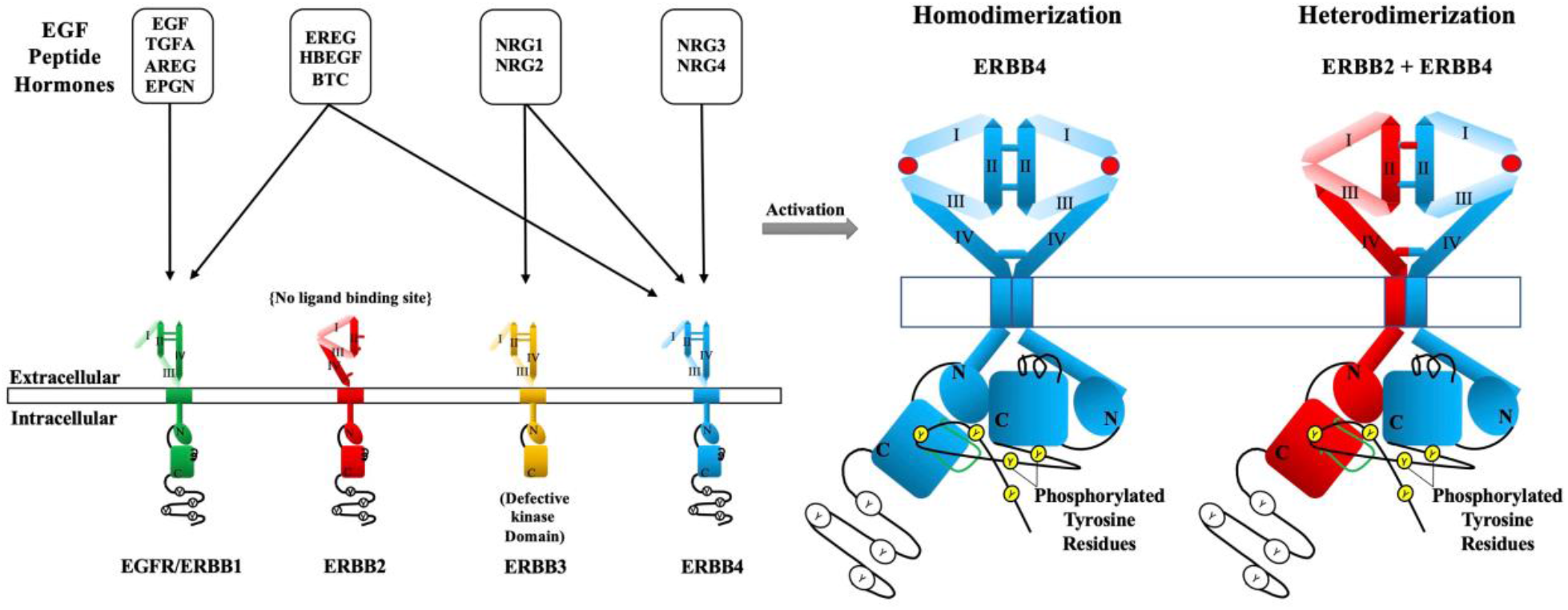
ERBB4 ligands stimulate ERBB receptor signaling via ERBB4 homodimerization and heterodimerization. Adapted from [4, 5].

ERBB4, like the other ERBB receptors, can undergo both homodimerization and heterodimerization [5] (**Figure 2**). Given that increased ERBB4 signaling can drive the proliferation of the IPC-298, MEL-JUSO, MeWo, and SK-MEL-2 *BRAF*-WT&*RAS*/*NF1-*mutant melanoma cell lines, we were somewhat surprised to observe that the synthetic, constitutively homodimerized and active Q646C mutant allele of *ERBB4* appears to potently <u>inhibit</u> the proliferation of the Mel-Juso, MeWo, and SK-MEL-2 *BRAF*-WT&*RAS*/*NF1-*mutant melanoma cell lines [4]. This apparent paradox may be rationalized by a rather simple observation. In many contexts, it appears that ERBB4 homodimers inhibit cell proliferation, whereas ERBB4-EGFR or ERBB4-ERBB2 heterodimers appear to stimulate cell proliferation [5].

To test this hypothesis, here we have determined the effects of wild-type *EGFR*, wild-type *ERBB2*, the K721A kinase-deficient (dominant-negative) *EGFR* mutant allele, and the K753A kinase-deficient (dominant-negative) *ERBB2* mutant allele on the clonogenic proliferation of the IPC-298, MEL-JUSO, MeWo, and SK-MEL-2 ERBB4-dependent, *BRAF*-WT&*RAS*/*NF1-*mutant melanoma cell lines.

## RESULTS

### A. ERBB2 is required for clonogenic proliferation of the IPC-298, MEL-JUSO, and MeWo BRAF-WT&RAS/NF1-mutant melanoma cell lines

We have infected the IPC-298, MEL-JUSO, MeWo, and SK-MEL-2 *ERBB4*-dependent, *BRAF*-WT&*RAS*/*NF1-*mutant melanoma cell lines with recombinant retroviruses based on pLXSN (which carries the neomycin resistance gene) to determine whether genes of interest stimulate or inhibit the formation of G418-resistant colonies of cells. This strategy is described in detail in the Materials and Methods section of this manuscript. A representative result is shown in **Figure 3** and complete, tabulated results are shown in **Table 1**.

**Table 1.** The kinase-deficient (dominant-negative) K753A *ERBB2* mutant allele inhibits the clonogenic proliferation of the IPC-298, MEL-JUSO, and MeWo melanoma cell lines. The kinase-deficient (dominant-negative) K721A *EGFR* mutant allele inhibits the clonogenic proliferation of the MeWo melanoma cell line. Neither the *ERBB2* K753A mutant allele nor the *EGFR* K721A mutant allele inhibits the clonogenic proliferation of the SK-MEL-2 melanoma cell line.

| Virus | IPC-298 (Female) |  |  |  | MEL-JUSO (Female) |  |  |
| --- | --- | --- | --- | --- | --- | --- | --- |
|  | Clonogenic Proliferation | Trials | P-Value |  | Clonogenic Proliferation | Trials | P-Value |
| LXSN | 100% | 4 |  |  | 100% | 6 |  |
| ERBB4 WT | 203% | 4 | 0.05 |  | 254% | 6 | 0.005 |
| EGFR WT | 307% | 4 | 0.08 |  | 266% | 6 | 0.001 |
| EGFR DN | 73% | 4 | 0.21 |  | 150% | 6 | 0.22 |
| ERBB2 WT | 171% | 4 | 0.02 |  | 235% | 6 | 0.003 |
| ERBB2 DN | 17% | 4 | 0.0001 |  | 52% | 6 | 0.02 |
| Virus | MeWo (Male) |  |  |  | SK-MEL-2 (Male) |  |  |
|  | Clonogenic Proliferation | Trials | P-Value |  | Clonogenic Proliferation | Trials | P-Value |
| LXSN | 100% | 6 |  |  | 100% | 5 |  |
| ERBB4 WT | 396% | 6 | 0.006 |  | 207% | 5 | 0.08 |
| EGFR WT | 162% | 6 | 0.05 |  | 605% | 5 | 0.03 |
| EGFR DN | 23% | 6 | 0.00002 |  | 523% | 5 | 0.08 |
| ERBB2 WT | 132% | 6 | 0.32 |  | 518% | 5 | 0.06 |
| ERBB2 DN | 19% | 6 | 0.000001 |  | 174% | 5 | 0.24 |

**Figure 3.**
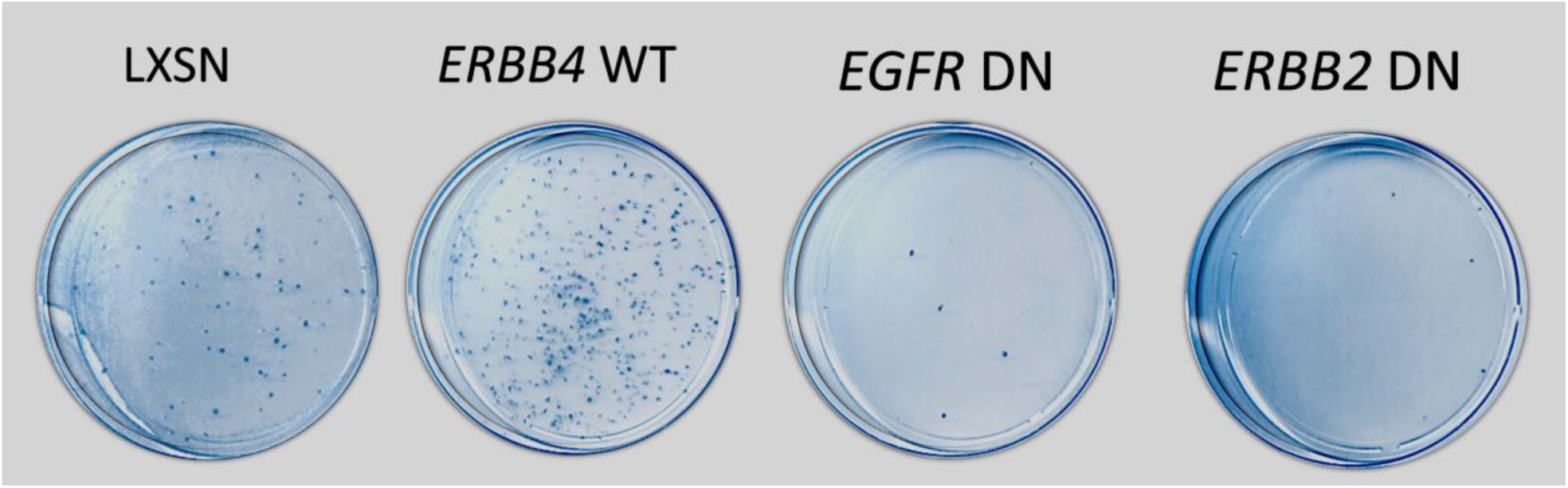
MeWo cells infected with recombinant retroviruses that express the K721A kinase-deficient (dominant-negative) *EGFR* mutant allele and the K753A kinase-deficient (dominant-negative) *ERBB2* mutant exhibit decreased clonogenic proliferation.

As shown in **Figure 3** and **Table 1**, wild-type *ERBB4* stimulates greater clonogenic proliferation than the LXSN vector control in the MeWo cell line. Wild-type *ERBB4* also stimulates greater clonogenic proliferation of the IPC-298, MEL-JUSO, and SK-MEL-2 cell lines (**Table 1**). These results are consistent with our previous observation that *ERBB4* is sufficient and necessary for the clonogenic proliferation of the MeWo, IPC-298, MEL-JUSO, and SK-MEL-2 cell lines.

As shown in **Figure 3** and **Table 1**, the kinase-deficient (dominant-negative) *ERBB2* K753A mutant allele causes less clonogenic proliferation than the LXSN vector control in the MeWo cell line. The *ERBB2* dominant-negative (DN) mutant allele also causes less clonogenic proliferation than the LXSN vector control in the IPC-298 and MEL-JUSO cell lines (**Table 1**). However, the *ERBB2* DN mutant allele does not cause less clonogenic proliferation than the LXSN vector control in the SK-MEL-2 cell line (**Table 1**). Thus, *ERBB2* appears to be necessary for the clonogenic proliferation of the MeWo, IPC-298, and MEL-JUSO cell lines, but does not appear to be necessary for the clonogenic proliferation of the SK-MEL-2 cell line.

### B. EGFR is required for clonogenic proliferation of the MeWo BRAF-WT&RAS/NF1-mutant melanoma cell line

As shown in **Figure 3** and **Table 1**, the kinase-deficient (dominant-negative) EGFR K721A mutant allele causes less clonogenic proliferation than the LXSN vector control in the MeWo cell line. However, the *EGFR* DN mutant allele does not cause less clonogenic proliferation than the LXSN vector control in the IPC-298, MEL-JUSO, and SK-MEL-2 cell lines (**Table 1**). Thus, *EGFR* appears to be necessary for the clonogenic proliferation of the MeWo cell lines, but does not appear to be necessary for clonogenic proliferation of the IPC-298, MEL-JUSO, or SK-MEL-2 cell line.

## MATERIALS AND METHODS

### Cell Lines and cell culture

Mouse C127 fibroblasts and the ψ2 and PA317 recombinant retrovirus packaging cell lines are generous gifts of Daniel DiMaio (Yale University). These cells were cultured essentially as described previously [6, 7]. The IPC-298 [8], MEL-JUSO [9], MeWo [10], and SK-MEL-2 [11] human melanoma cell lines were obtained from ATCC [12] (Manassas, Virginia, USA) and DSMZ [13] (Braunschweig, Germany). These cell lines were cultured as recommended or as previously described [7-11]. Cell culture media, serum, and supplements were obtained from Cytiva [14] (Marlborough, VA). G418 was obtained from Corning [15] (Corning, NY).

### Recombinant retrovirus constructs

The recombinant retroviral expression constructs pLXSN, pLXSN-ERBB4, pLXSN-EGFR, and pLXSN-ERBB2 possess the neomycin-resistance gene and have been described previously [16, 17]. The recombinant retroviral expression constructs pLXSN-EGFR-K721A and pLXSN-ERBB2-K753A have been described previously [18].

### Packaging recombinant ecotropic and amphotropic retroviruses

Recombinant amphotropic retroviruses were packaged using the ψ2 ecotropic retrovirus packaging cell line and the PA317 amphotropic retrovirus packaging cell line as previously described [16, 19, 20]. Briefly, the recombinant retrovirus plasmids were stably transfected using calcium phosphate into the ψ2 ecotropic retrovirus packaging cell line. Transfected cells were selected using G418. Colonies of G418-resistant cells were pooled to create a stable cell line for each transfectant. Low-titer (∼10^4^ infectious units/mL) ecotropic retrovirus stocks were generated by harvesting the conditioned medium from each stably-transfected ψ2 cell line. PA317 cells were infected with the low-titer ecotropic retrovirus stocks. Colonies of G418-resistant cells were pooled to create a stable cell line for each infection. High-titer (∼10^6^ infectious units/mL) amphotropic retrovirus stocks were generated by harvesting the conditioned medium from each stably-infected PA317 cell line.

### Clonogenic proliferation assay

Clonogenic proliferation assays were performed essentially as described [1, 4, 19, 21-24]. A 60 mm dish each of the C127 cell line and one or more tester cell lines (IPC-298, MEL-JUSO, MeWo, and SK-MEL-2) were infected with the LXSN vector control retrovirus and the experimental retroviruses (LXSN-ERBB4, LXSN-EGFR, LXSN-EGFR-K721A, LXSN-ERBB2, and LXSN-ERBB2-K753A). Each 60 mm dish of cells was subcultured into three 100 mm dishes and the infected cells were selected using G418. The resulting colonies of G418-resistant cells were stained using Giemsa. The stained plates were digitized and the drug-resistant colonies were counted. The titer of each retrovirus in each cell line was determined by dividing the total number of drug-resistant colonies (in the three 100 mm dishes) of the infection by the volume of retrovirus used for the infection. To calculate the efficiency of clonogenic proliferation, the titers of each retrovirus in the tester cell lines was divided by the titer of the same retrovirus in the C127 cells. In the tester cell lines, the efficiency of clonogenic proliferation for each of the experimental retrovirus was normalized against the efficiency of clonogenic proliferation for the LXSN retrovirus. The normalized efficiency of clonogenic proliferation was averaged from multiple trials. If the average normalized efficiency of clonogenic proliferation is significantly greater (one sample t test) than 100%, then the gene of interest contained in that retrovirus is judged to have stimulated clonogenic proliferation. Likewise, if the average normalized efficiency of clonogenic proliferation is significantly less (one sample t test) than 100%. Then the gene of interest contained in that retrovirus is judged to have inhibited clonogenic proliferation. A P value of 0.05 (1-tailed) was used as a threshold for significance.

## DISCUSSION

### A. ERBB4-ERBB2 heterodimers appear to drive three (IPC-298, MEL-JUSO, and MeWo) ERBB4-dependent, BRAF-WT&RAS/NF1-mutant melanoma cell lines

We have previously observation that the synthetic, constitutively dimerized and active *ERBB4* Q646C mutant allele inhibits the clonogenic proliferation of the *ERBB4*-dependent MEL-JUSO, MeWo, and SK-MEL-2 cell lines [4]. Here we show that the *ERBB2* DN mutant allele inhibits the clonogenic proliferation of the *ERBB4*-dependent IPC-298, MEL-JUSO, and MeWo cell lines. This combination of observations suggests that ERBB4-ERBB2 heterodimers are driving the proliferation of IPC-298, MEL-JUSO, and MeWo cell lines. Because there are numerous approved inhibitors of ERBB2 signaling, including one agent (pertuzumab) that inhibits signaling by ERBB2 heterodimers [25-28], the translational implications are obvious.

### B. ERBB4 appears to possess gender-selective effects in vivo

Our *in silico* analysis of deceased *BRAF* WT melanoma patients indicates that the deceased male patients are much more likely to be associated with an *ERBB4* mutant allele than the deceased female patients [4]. This suggests that increased ERBB4 signaling is more relevant to the genesis and aggressiveness of melanoma in males than in females. Indeed, the gender-selective effects of *ERBB4* may account for the fact that *BRAF* WT melanomas are more deadly in males than in females [1, 4].

Here we observe that the female IPC-298 and MEL-JUSO cell lines are dependent on *ERBB2*, whereas the male MeWo cell line is dependent on both *EGFR* and *ERBB2*. This difference suggests that the combination of ERBB4-ERBB2 and ERBB4-EGFR heterodimers make *BRAF* WT melanomas more deadly in males than in females. One possibility is that EGFR potentiates the action of testosterone on ERBB4 via the cytoplasmic LXXLL motif found in ERBB4, which regulates interactions of ERBB4 with steroid hormone receptor co-activators [5].

It is also noteworthy that the male SK-MEL-2 cell line is not dependent on *EGFR* nor *ERBB2*. The obvious conclusion that neither ERBB4-EGFR nor ERBB4-ERBB2 heterodimers drive SK-MEL-2 cells is at least superficially contrary to our general model for ERBB4 function [5]. However, ERBB4 homodimers encoded by the non-canonical *ERBB4* JM-a/CT-b splicing isoform drive proliferation in a variety of contexts [5, 29], suggesting that SK-MEL-2 cells express the *ERBB4* JM-a/CT-b splicing isoform rather than the canonical *ERBB4* JM-a/CT-a isoform, thereby enabling ERBB4 homodimers to drive the proliferation of these cells.

